# Emerging chikungunya virus variants at the E1-E1 inter-glycoprotein spike interface impact virus attachment and Inflammation

**DOI:** 10.1101/2021.09.13.460192

**Authors:** Margarita V. Rangel, Nicole McAllister, Kristen Dancel-Manning, Maria G. Noval, Laurie A. Silva, Kenneth A. Stapleford

**Affiliations:** Department of Microbiology, New York University Grossman School of Medicine, New York, New York 10016; Department of Biology, Seton Hill University, 1 Seton Hill Drive, Greensburg, Pennsylvania 15601; Microscopy Laboratory of Division of Advanced Research Technologies, New York University Langone Medical Center, New York, New York 10016; Department of Microbiology and Molecular Genetics, University of Pittsburgh School of Medicine, Pittsburgh, Pennsylvania 15261; Department of Pediatrics, University of Pittsburgh School of Medicine, Pittsburgh, Pennsylvania 15261

## Abstract

Chikungunya virus (CHIKV) is a re-emerging arthropod-borne alphavirus and a serious threat to human health. Therefore, efforts toward elucidating how this virus causes disease and the molecular mechanisms underlying steps of the viral replication cycle are crucial. Using an *in vivo* transmission system that allows intra-host evolution, we identified an emerging CHIKV variant carrying a mutation in the E1 glycoprotein (V156A) in the serum of mice and saliva of mosquitoes. E1 V156A has since emerged in humans during an outbreak in Brazil, co-occurring with a second mutation, E1 K211T, suggesting an important role for these residues in CHIKV biology. Given the emergence of these variants, we hypothesized that they function to promote CHIKV infectivity and subsequent disease. Here, we show that E1 V156A and E1 K211T modulate virus attachment and fusion and impact binding to heparin, a homolog of heparan sulfate, a key entry factor on host cells. These variants also exhibit differential neutralization by anti-glycoprotein monoclonal antibodies, suggesting structural impacts on the particle that may be responsible for altered interactions at the host membrane. Finally, E1 V156A and E1 K211T exhibit increased titers in an adult arthritic mouse model and induce increased foot-swelling at the site of injection. Taken together, this work has revealed new roles for E1 where discrete regions of the glycoprotein are able to modulate cell attachment and swelling within the host.

**IMPORTANCE:** Alphaviruses represent a growing threat to human health worldwide. Chikungunya virus (CHIKV) has rapidly spread to new geographic regions in the last several decades, causing overwhelming outbreaks of disease, yet there are no approved therapeutics. The CHIKV glycoproteins are key determinants of CHIKV adaptation and virulence. In this study, we characterize the naturally emerging E1 glycoprotein variants, V156A and K211T. We demonstrate that E1 V156A and K211T function in virus attachment to cells, a role that until now has been only attributed to the CHIKV E2 glycoprotein. We also demonstrate E1 V156A and K211T to increase foot-swelling in mice. Observing that these variants and other pathogenic variants occur at the E1-E1 inter-spike interface, we highlight this structurally important region as critical for multiple steps during CHIKV infection. Together, these studies further defines the function of E1 in CHIKV infection and can inform the development of therapeutic or preventative strategies.

## INTRODUCTION

Arboviruses are an expansive group of human pathogens that constitute a continuing global health threat. Over the last several decades, there has been a resurgence of arbovirus outbreaks with high morbidity and mortality, demonstrating the ability of arboviruses to adapt and spread to new environments [1]. Alphaviruses represent a prominent threat as members of this family of arboviruses, including chikungunya virus (CHIKV), eastern equine encephalitis virus, O’nyong’nyong virus, Sindbis virus, and Mayaro virus and have led to outbreaks in Asia, Europe, Africa, and the Americas [2–5]. To date there are no vaccine or antiviral therapies targeting alphaviruses, therefore a better understanding of the mechanisms underlying how these viruses cause disease is crucial to identifying new therapeutic targets.

CHIKV is a re-emerging pathogen that has caused explosive outbreaks and spread throughout the world including the Americas [2, 6]. In 2020 alone, there were large outbreaks reported around the world, each consisting of 30,000-100,000 cases as well as smaller outbreaks of hundreds of cases [7]. CHIKV has a single-stranded positive-sense RNA genome that encodes two open reading frames (ORFs). The first ORF encodes four non-structural proteins (nsP1-4) and the second encodes six structural proteins (capsid, E3, E2, 6K, TF, and E1) [8]. The mature CHIKV particle consists of 240 copies of the E1 and E2 glycoproteins, the fusion protein and attachment protein, respectively. The glycoproteins are arranged in trimeric spikes (E1-E2)_3_ of heterodimers in T=4 icosahedral symmetry with E1-E2 inter-dimer and E1-E1 inter-spike contacts stabilizing the protein lattice [9]. The glycoprotein spikes protrude from a host-derived lipid bilayer that surrounds the nucleocapsid core [10] containing the RNA genome.

It has been demonstrated that CHIKV can engage various host cell factors and putative receptors such as glycosaminoglycans (GAGs), C-type lectins, and the recently identified receptor Mxra8 [11–13]. Thus far, specific virus-host interactions have been attributed to the E2 glycoprotein. For instance, binding to the ubiquitous host GAGs is, in part, modulated by the type of amino acid at E2 residue 82 [11, 14]. Residue 82 is a highly conserved glycine in nearly every CHIKV isolate, except for CHIKV 181/25, which has an arginine at residue 82 and altered GAG utilization [14–16]. Moreover, the CHIKV 181/25 E2 residue D71 has been shown to be critical for interactions with Mxra8 [17]. However, interestingly Mxra8 is predicted to contact residues within both E1 and E2, suggesting both proteins of the spike have the potential to impact cell attachment [18].

Following attachment, the particle is taken up by clatherin-mediated endocytosis and pH-dependent membrane fusion occurs in the early endosome [19, 20]. Upon exposure to low pH, the E1-E2 heterodimer dissociates, exposing the fusion peptide [21]. The fusion peptide, consisting of residues 83-100 of the E1 *cd* loop, inserts into the target membrane and E1 undergoes a conformational change, folding back on itself, and trimerizing [22–24]. The insertion of multiple E1 trimers mediates the joining of viral and host membranes [24, 25]. In addition to virus entry, the CHIKV glycoproteins play roles in egress of mature particles from infected cells, particle stability, and immunogenicity [24, 26, 27]. Considering this multi-functionality throughout the viral replication cycle, it is likely that there are additional roles for E1 and further details of its known functions yet to be elucidated.

The evolution and spread of CHIKV has been marked with mutations in the E1 and E2 envelope glycoproteins, making these proteins key determinants of infectivity and pathogenesis and pivotal for adaptation [28–30]. Following a 2005 outbreak, it was retrospectively observed that CHIKV acquired a single mutation in E1, A226V, that increased infectivity in an alternative vector, *Aedes (Ae.) albopictus*, as opposed to the primary vector, *Ae. aegypti*. This adaptation event led to outbreaks in areas of naïve populations where *Ae. albopictus* was abundant, giving rise to one of the four recognized lineages of CHIKV, the Indian Ocean Lineage (IOL) [31]. The IOL strain was also responsible for an explosive outbreak in South East Asia in 2008, where the endemic Asian Lineage had already long been circulating [32]. In questioning why the Asian Lineage had not yet gained E1 A226V despite the high abundance of *Ae. albopictus*, it was found that E1 98T of the Asian Lineage versus E1 98A of IOL has an epistatic interaction with E1 226, limiting the penetrance of E1 A226V mutation and, further, that E1 98A enhances penetrance [33]. These findings demonstrate the impact of adaptive CHIKV glycoprotein variants. Continued work to understand how the glycoproteins adapt to modulate CHIKV infection will aid in identifying ways to therapeutically target these critical viral proteins.

In this study, we identify the novel CHIKV E1 variant V156A in our mouse-mosquito transmission system that notably arose in humans with a second mutation K211T [34]. We provide a functional characterization of these residues to guide our developing knowledge of how E1 functions throughout the CHIKV replication cycle. We show that CHIKV E1 V156A and K211T, present at the E1-E1 inter-spike interface, influence cell binding and fusion and interactions with the GAG heparin. Moreover, we show that these phenotypes are possibly driven by a structural change in the glycoproteins, as suggested by altered neutralization by both E1 and E2 targeting monoclonal antibodies. Finally, we demonstrate CHIKV E1 V156A and K211T lead to increased titers and foot-swelling in a mouse model, with the latter being the first observation of a discrete E1 residue to have this impact. Together, our findings highlight new roles for E1 in modulating binding to cells and pathogenesis in animals and identifies E1 residues 156 and 211 as key determinants of virulence.

## RESULTS

### Emergence of epidemic chikungunya virus E1 variant V156A in mice

In a previous study to understand chikungunya virus (CHIKV) evolution during vector-to-vertebrate host transmission, we infected *Ae. aegypti* mosquitoes with an IOL strain of CHIKV via artificial bloodmeal and allowed these mosquitoes to feed on neonatal mice [35]. We then analyzed the emerging virus populations by deep sequencing and identified several novel mutations in the E1 glycoprotein (V80I and A129V) that increase replication and transmission *in vivo* [28, 35]. In addition to these variants, we also identified another emerging mutation in the E1 glycoprotein at residue 156 (V156A) (Figure 1). E1 V156A could be detected at low frequency, yet above background (>0.01%) in the saliva of some bloodmeal-infected mosquitoes. Importantly, in subsequent mice and mosquitoes, E1 V156A and was able to increase in frequency, displacing the parental virus in some cases (Figure 1A and B). Although this result was variable between transmission events, when we performed an additional round of transmission from mouse-fed mosquito to mouse, V156A remained fixed in both mosquito and mouse populations (Figure 1C). Significantly, several years after our transmission study, CHIKV E1 V156A was detected in the serum of infected humans during an outbreak in Brazil [34]. Interestingly, whereas we identified V156A in the IOL background, it appeared in nature in the East Central South African (ECSA) background and co-occurred with the E1 mutation K211T each time it was detected [34].

**Figure 1:**
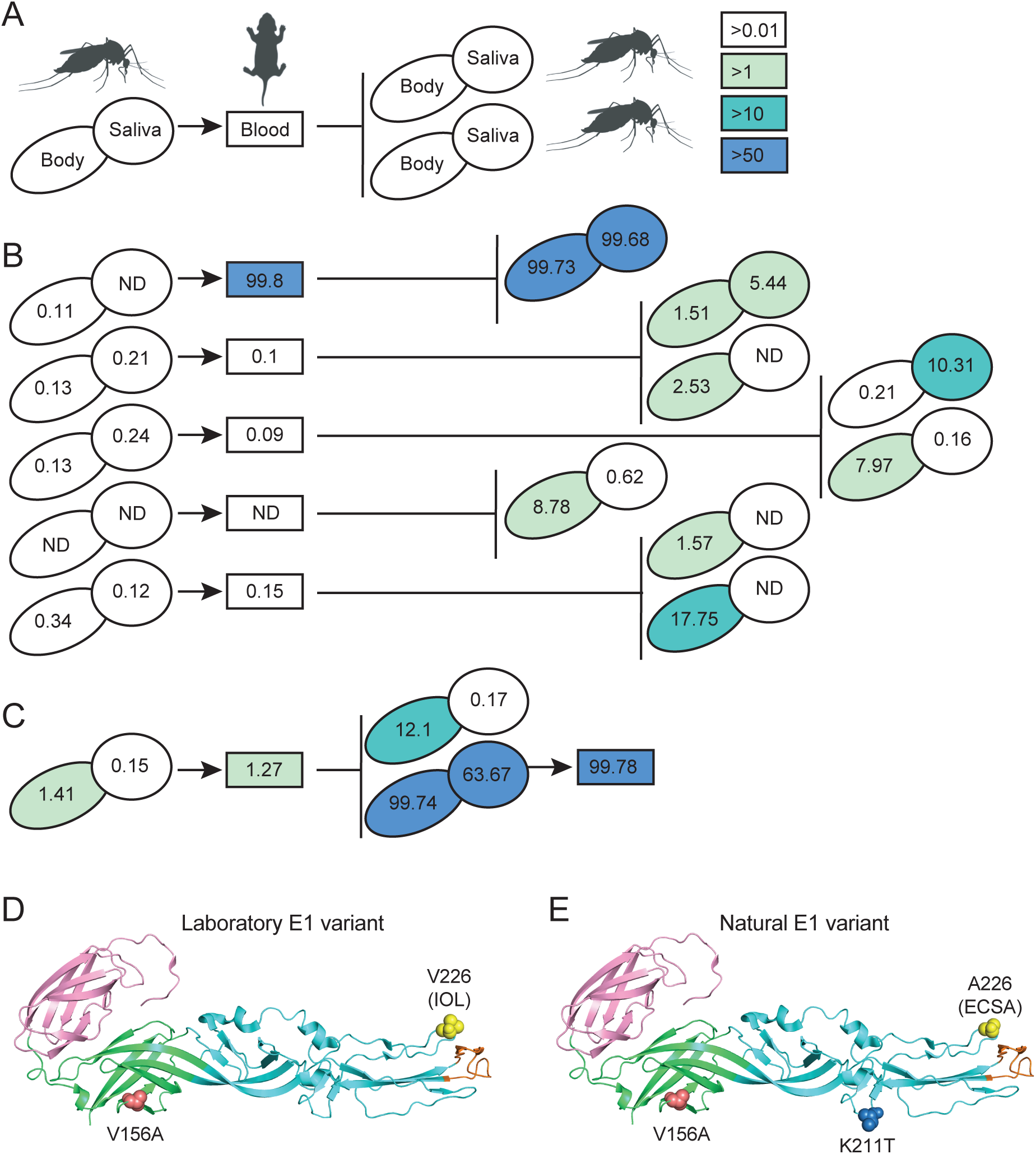
Identification of emerging CHIKV E1 variant V156A following vector-borne transmission. (A) Schematic representation of vector-borne transmission system and color key for mutation frequency detected in mosquito bodies or saliva (ovals) and mouse blood (rectangles). ND = not detected, below background (<0.01%). (B) Frequency of V156A detection during single rounds of vector-borne transmission, color-coded as depicted in (A). (C) Frequency of V156A detection during two rounds of vector-borne transmission, color-coded as depicted in (A). (D) Structure of E1 monomer depicting the laboratory-identified E1 variant with residues 156 and 226 indicated. E1 domain I is colored green, domain II cyan, and domain III pink. E1 fusion loop is colored orange. (E) Structure of E1 monomer depicting the E1 variant identified in nature with residues 156, 211 and 226 indicated. The background limit of detection was <0.01%.

Positions 156 and 211 are located on the same face of the E1 glycoprotein, in domains I and II, respectively (Figure 1D and E). Position 156 is in the hinge region of E1 and near the domain I-domain III linker, a dynamic region important for the conformational change of E1 that contributes to the facilitation of membrane fusion (Figure 1D, red) [36]. Position 211 is in a region of domain II distal to the fusion loop, a region containing inter-trimer interacting regions between post-fusion E1 trimers (Figure 1E, blue) [24]. On the mature particle, position 156 and 211 of adjacent E1 monomers face each other at the E1-E1 inter-spike interface. The IOL lineage originated from the ECSA lineage and carries the epidemic E1 A226V mutation (Figure 1D and E, yellow) [6]. It is possible that these evolving E1 variants represent discrete residues critical for protein function. Considering their emergence in our transmission system and in nature, we hypothesized that E1 V156A and E1 K211T function to provide a replicative advantage during CHIKV infection.

### CHIKV E1 V156A and K211T exhibit replication kinetics and infectious particle production similar to wild-type CHIKV *in vitro*

To begin characterizing the E1 variants, we generated both single variants (IOL-V156A and IOL-K211T) and a double variant (IOL-V156A:K211T) on the IOL background as well as single and double variants on a version of the IOL background with residue V226 mutated to A226 (V226A) (Figure 1D and E). We first performed multi-step replication curves in BHK-21 cells to assess the replication of each variant (Figure 2A and B). We found that infectious titers of the variants exhibited a statistically significant decrease, although this was to a modest extent and overall, we consider them comparable to wild-type CHIKV (Figure 2A and B). Notably, by 48 hours the titers of the variants on the 226V background were significantly lower than the wildtype control when compared with the titers of the counterpart variants on the 226A background, suggesting that 226A may be providing an advantage for the impaired titers of V156A and K211T. We then inspected the CHIKV structural proteins in purified stocks of each virus by western blot. We found comparable levels in E1, E2, and capsid in each virus (Figure 2C and D). Finally, to investigate any morphological differences between variants, we visualized infectious mature particles via transmission electron microscopy (TEM). We found each virus to produce particles of similar size and uniformity (Figure 2E). These results demonstrate that E1 V156A and E1 K211T variants have similar, growth kinetics and infectious particle production to wild-type CHIKV.

**Figure 2:**
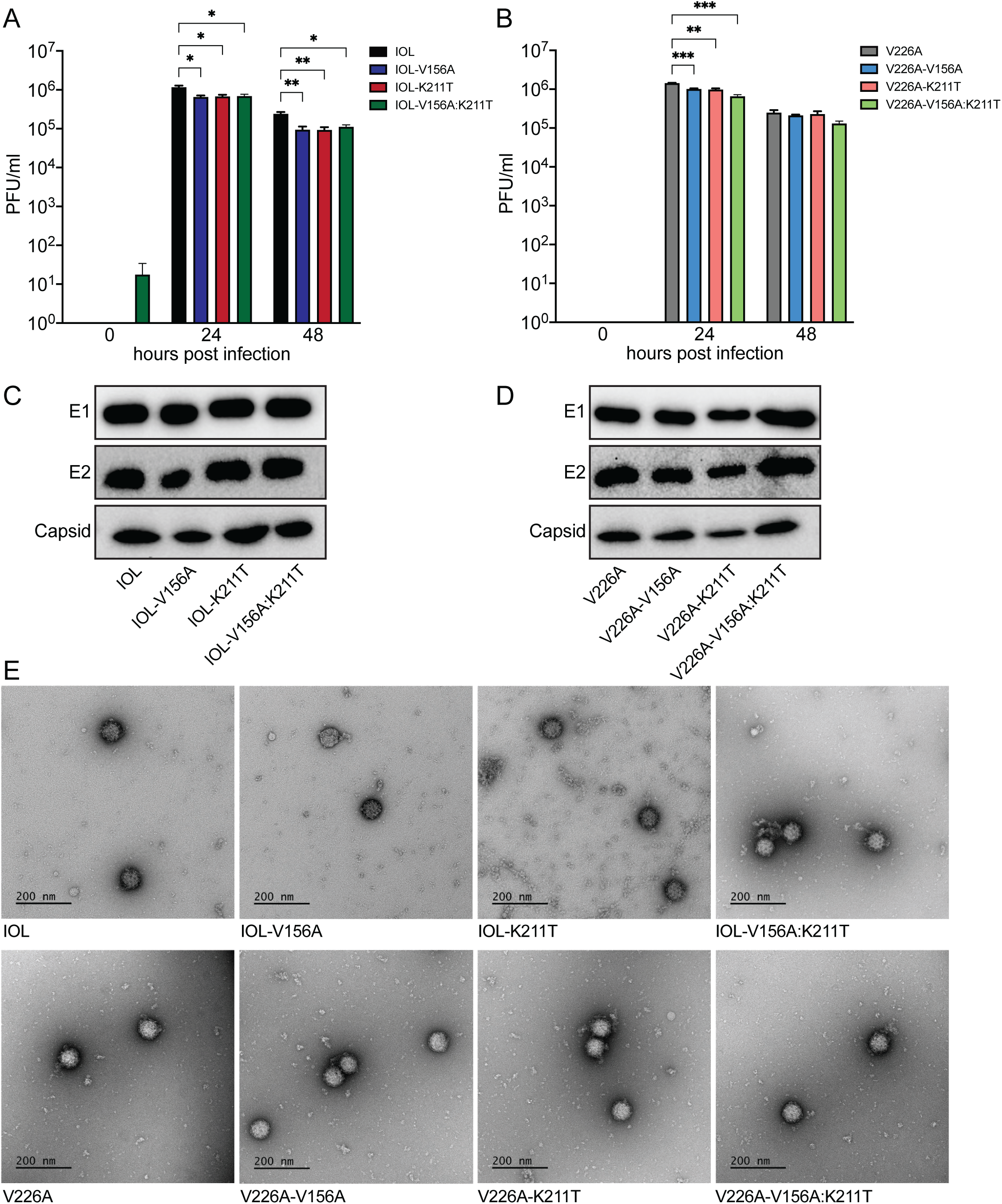
Replication kinetics and particle production of E1 variants resemble wild-type CHIKV. (A and B) Multi-step replication curves for wild-type CHIKV and E1 variants in BHK-21 cells. BHK-21 cells were infected at an MOI of 0.1 PFU/cell and infectious titers of supernatant fractions collected at each timepoint were determined by plaque assay. Data represents two independent experiments performed in triplicate, * p<0.05, ** p<0.01 and *** p<0.001. P values were determined by two-way ANOVA with Dunnett’s multiple comparison test. (C and D) Immunoblotting of structural proteins of purified particles of wild-type CHIKV or E1 variants. Equivalent amounts of infectious particles were suspended in Laemmli buffer and proteins were separated by SDS-PAGE and analyzed by immuoblotting. Blots represent one of two independent viral stocks. (E) TEM imaging of purified particles of CHIKV wild-type and E1 variants.

### CHIKV E1 V156A and K211T promote entry and exhibit increased sensitivity to changes in endosomal pH

The primary recognized role of E1 is the facilitation of low pH-triggered membrane fusion, a necessary step in establishing infection for enveloped viruses. Given the location of residues 156 and 211 in regions important for mediating fusion, we determined the capacity of each of the variants capacity to undergo membrane fusion at various pHs. To test this, we performed a fusion-from-without assay on BHK-21 cells (Figure 3A and B). We compared each variant with its parental virus, IOL or V226A, and as a control, included the E1 variant V80L, which we have demonstrated to have a decreased pH threshold for fusion [28]. We observed that, on the IOL background, while the single variants exhibited a similar pH threshold for fusion as the parental virus, the double mutant IOL-V156A:K211T consistently resulted in an increase in the percentage of infected cells at low pH values (Figure 3A). Interestingly, this was not the case on the V226A background in which all V226A variants, including the double mutant V226A-V156A:K211T, resembled the parental virus (Figure 3B). These results suggest that residues 156, 211, and 226 function together to promote fusion or provide an entry advantage such as increased cell attachment.

**Figure 3:**
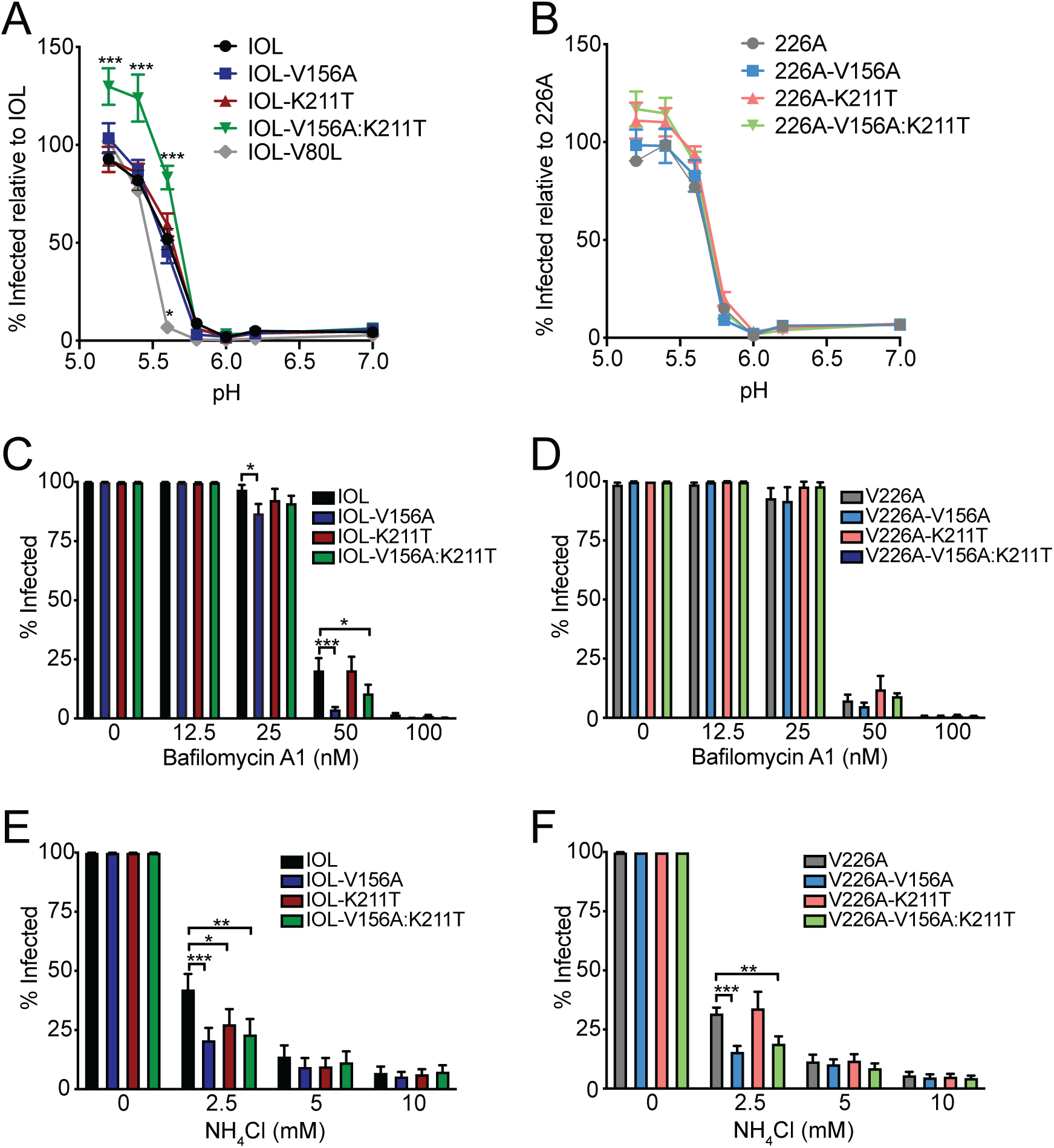
CHIKV E1-V156A and K211T variants modulate cell entry and increase sensitivity to endosomal pH. (A and B) Fusion-from-without assay. BHK-21 cells were preincubated with NH_4_Cl, adsorbed with ZsGreen-expressing viruses (MOI=1), fusion was triggered for 2 min at the indicated range of pH, and viral spread was blocked by replenishing NH_4_Cl-containing media. 18 h post infection, cells were fixed, DAPI stained, and the percent of infected cells was assessed by fluorescence imaging. Bafilomycin A1 (C and D) and NH_4_Cl (E and F) sensitivity assays. BHK-21 cells were preincubated with Bafilomycin A1 or NH_4_Cl, incubated with ZsGreen-expressing virus (MOI=1) for 1 h, washed twice, and incubated in media containing each treatment for 18 h. Cells were then fixed, DAPI stained, and the percent of infected cells was assessed by fluorescence imaging. All data is representative of at least three independent experiments performed in duplicate, * p<0.05, ** p<0.01 and *** p<0.001. P values were determined by two-way ANOVA with Bonferroni’s multiple comparison test.

As a complementary approach to assess the pH-dependence of the E1 variants, we tested their infectivity on BHK-21 cells in the presence of NH_4_Cl or Bafilomycin A1, lysosomotropic agents used to deacidify the endosome [37–39]. IOL-V156A and IOL-V156A:K211T exhibited increased sensitivity to Bafilomycin A1 (Figure 3C), yet this was not observed for their V226A counterparts (Figure 3D). V156A and V156A:K211T on both IOL and V226A backgrounds were more sensitive to NH_4_Cl whereas the K211T single mutant was more sensitive to NH_4_Cl on the IOL background but not the V226A background (Figure 3E and F). The varying sensitivity of the variants to lysosomotropic agents further suggests unique entry dynamics dictated by E1 residues 156, 211, and 226.

### CHIKV E1 V156A and K211T decrease cell binding

One possible explanation for the increased infectivity exhibited by IOL-V156A:K211T in our fusion-from-without assay (Figure 3A) may be due to an enhanced binding of this variant to cells. To test this hypothesis, we conducted binding assays on mammalian and mosquito cell types (Figure 4). To differentiate the impact on binding by V156A and K211T apart from E1 V226A, we compared binding of wild-type CHIKV E1 V226 and E1 A226 (Figure 4A). Interestingly, we found A226 exhibits decreased binding on BHK-21 and both insect cell lines, suggesting that E1 residue 226 influences cell binding. We then investigated the binding of each E1 variant. In BHK-21 cells and both insect cell lines, we found that the single and double variants on the IOL (V226) background significantly decrease cell binding compared with wild-type CHIKV (Figure 4B). V156A and K211T in the 226A background also impaired binding compared with the wildtype counterpart (Figure 4C). Taken together, these results suggest that the increased infectivity observed in the fusion assay for IOL-V156A:K211T (Figure 3A) cannot be explained by an increase in binding as binding of this variant to BHK-21 cells, as well as both insect cell lines, was decreased compared with wild-type CHIKV. The decrease in binding of these E1 variants in mammalian and insect cells suggest that these E1 specific residues may influence interactions with a host membrane component present on the surface of each of these cell types.

**Figure 4:**
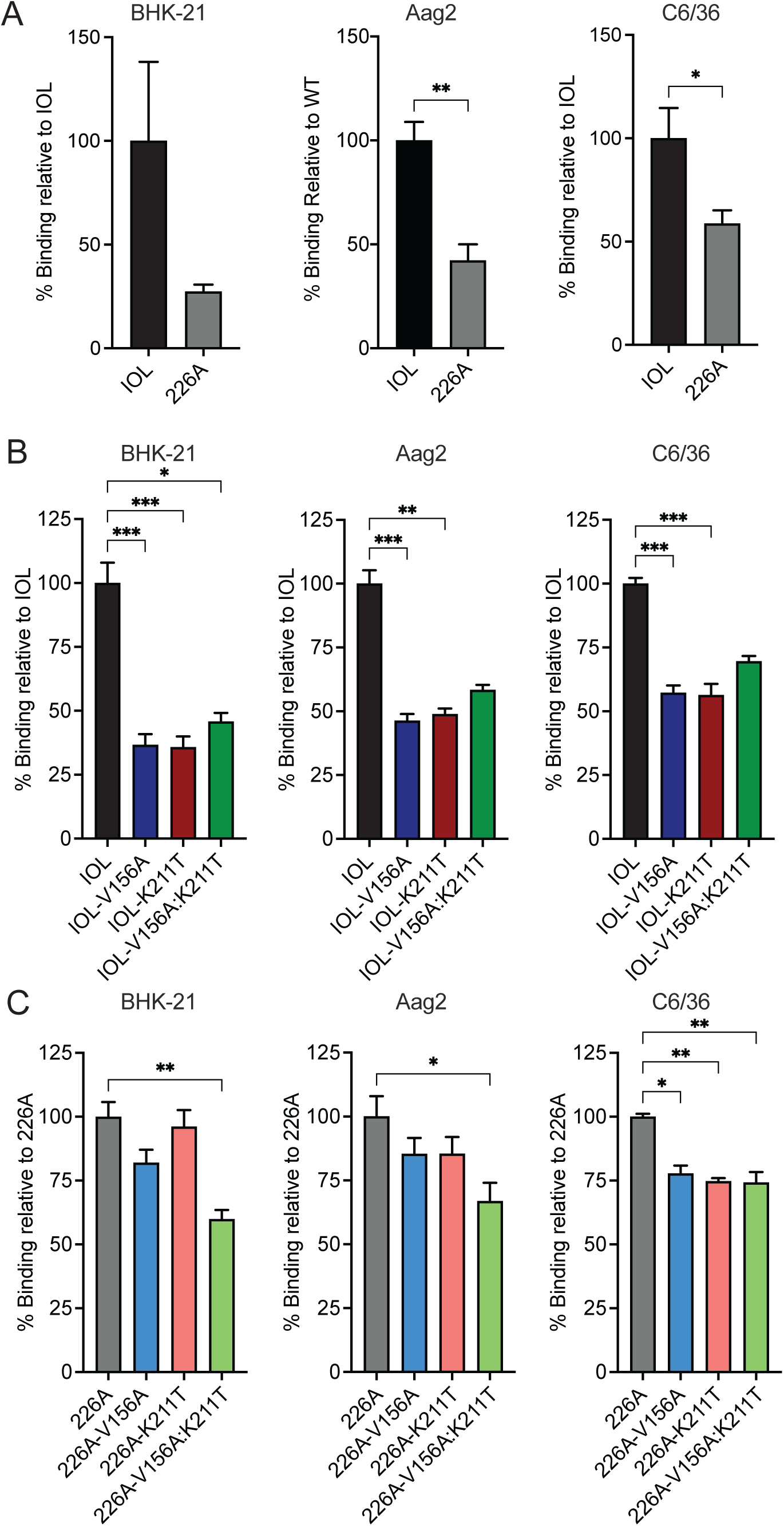
CHIKV E1 variants V156A and K211T exhibit impaired cell binding. Virus-cell binding assays comparing IOL and 226A viruses (A), V156A and K211T variants on the IOL background (B), and V156A and K211T variants on the 226A background (C). BHK-21, Aag2, or C6/36 cells were preincubated with NH_4_Cl at 4°C then adsorbed with viruses (10 genomes/cell) for 30 minutes. Cells were washed extensively and harvested for RNA extraction and cDNA synthesis. Percent relative binding was determined using SYBR Green qPCR with primers targeting CHIKV or cellular GAPDH (BHK-21) or actin (Aag2 and C6/36). All data is representative of at least three independent experiments performed in duplicate, * p<0.05, ** p<0.01 and *** p<0.001. P values were determined by Mann-Whitney test (A) or Kruskal-Wallis test (B and C).

### CHIKV E1 V156A and K211T reduce CHIKV-heparin interactions

Considering the decreased binding across all cell types tested, we hypothesized that the E1 variants may have altered interactions with the ubiquitously expressed cell surface factors, GAGs, known to function as a CHIKV attachment factors [16, 40, 41]. To test this hypothesis, we measured the relative binding of each CHIKV E1 variant to heparin, a highly sulfated GAG similar in structure to heparan sulfate, using an ELISA. We found that the single variants IOL-V156A, IOL-K211T, and the double mutant all exhibited decreased binding to heparin (Figure 5A and B). Interestingly, binding of the individual variants in the V226A background was not as dramatically decreased. However, we did find a significant reduction in heparin binding with the V226A double variant (V226A-V156A:K211T). Together these results demonstrate residues 156, 211, and 226 have the potential to modulate interactions at the cell surface, particularly with important entry factors such as glycosaminoglycans.

**Figure 5:**
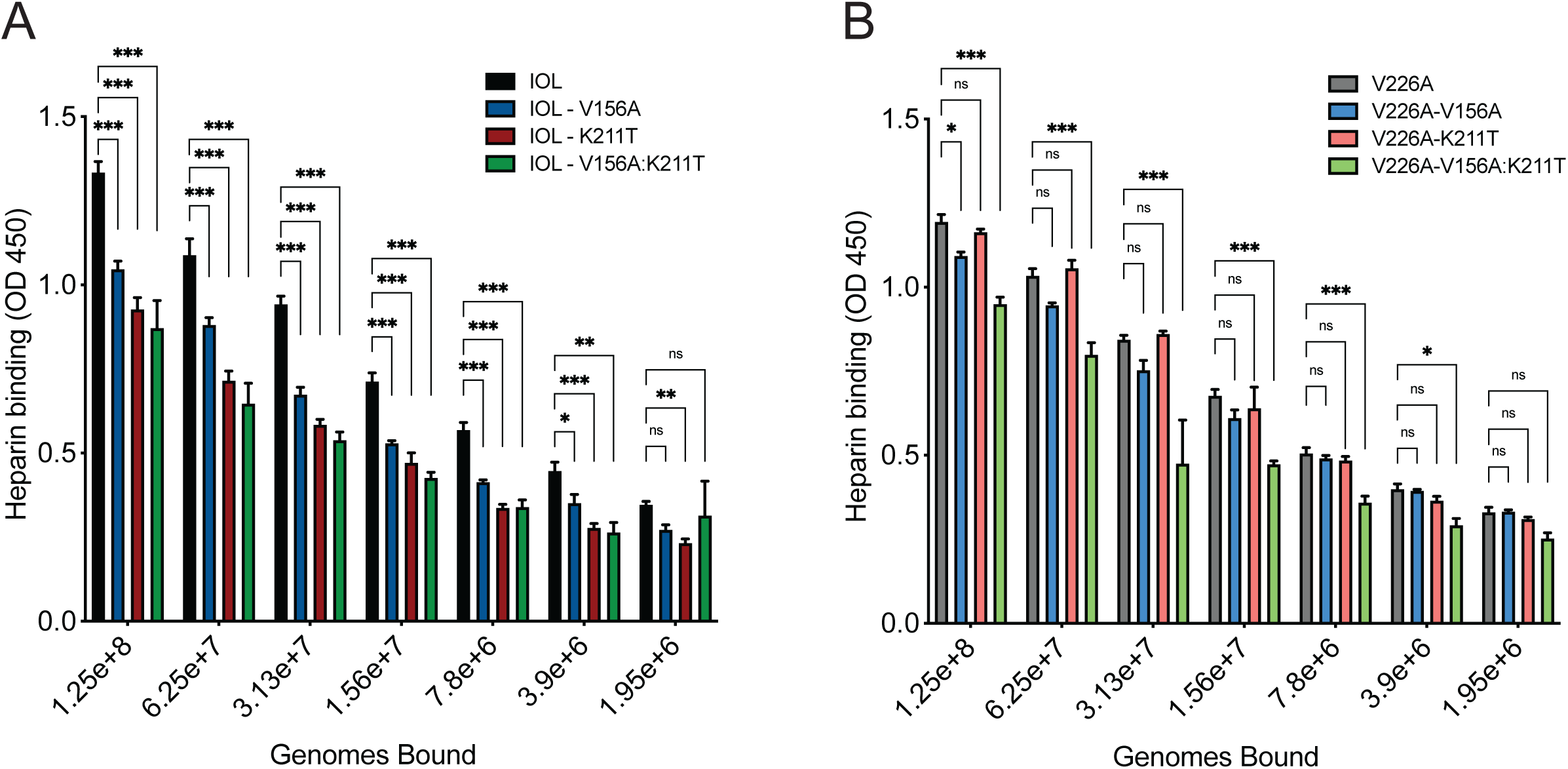
CHIKV E1 V156A and K211T decrease binding to heparin. Avidin-coated ELISA plates were bound with biotinylated heparin and incubated with serially-diluted virus of known genome number for 1-2 hours. Plates were washed to remove unbound virus. Bound virus was detected using mouse anti-CHIKV E2 mAb CHIK-187 followed by goat anti-mouse HRP-conjugated secondary antibody. Plates were developed with TMB substrate and absorbance was measured at 450 nm. Data is representative of two experiments performed in duplicate, * p<0.05, ** p<0.01 and *** p<0.001. P values were determined by two-way ANOVA with Bonferroni’s multiple comparison test.

### CHIKV E1 V156A and K211T impact antibody neutralization

We hypothesized that a mechanism by which E1 V156A and K211T may be leading to altered binding to cells is by conferring structural changes in the glycoproteins. To investigate whether E1 V156A and K211T induce local structural changes in the glycoprotein complex, we treated gradient-purified virions with increasing concentrations of murine monoclonal antibodies (mAbs) targeting different regions of E1 or E2 in neutralization assays (Figure 6A) [42]. Mapping of escape mutants revealed two E2 antibodies, CHIK-102 and CHIK-263, target the most distal region of E2, domain B, and E1 antibody CHIK-166 targets part of domain II of E1, near the shielded fusion loop (Figure 6A) [42]. CHIK-263 Fab bound to CHIKV was further analyzed by Cryo-EM and was found to have an epitope footprint that spans both E1 and E2 (Figure 6A) [43]. We hypothesized that changes in neutralization by these antibodies would suggest altered epitope orientation. We found that IOL-K211T and IOL-V156A:K211T were both more sensitive than wild-type CHIKV to neutralization by CHIK-102 and CHIK-263, while IOL-V156A was largely unaffected (Figure 6B and C). In addition, we found that IOL-K211T and IOL-V156:K211T were neutralized similarly by CHIK-263 whereas the double mutant was more sensitive than IOL-K211T to neutralization by CHIK-102. Finally, we found that IOL-V156A was more sensitive to neutralization by CHIK-166, than with the E2 antibodies (Figure 6D). IOL-K211T and IOL-V156A:K211T were less sensitive to neutralization by CHIK-166 than by CHIK-102 or CHIK-263, but still were more sensitive than IOL-V156A and wild-type. As a control, none of the variants were neutralized by an anti-ZIKV antibody (Figure 6E). Individual IC_50_ values are listed in Figure 6, panel F. The differential neutralization of the E1 variants compared to wild-type by anti-glycoprotein mAbs suggests that the variants V156A and K211T and cause structural changes in the particle leading to changes in the mAb epitopes. We hypothesize that these changes may also mediate the altered binding to cells and increased fusion observed.

**Figure 6:**
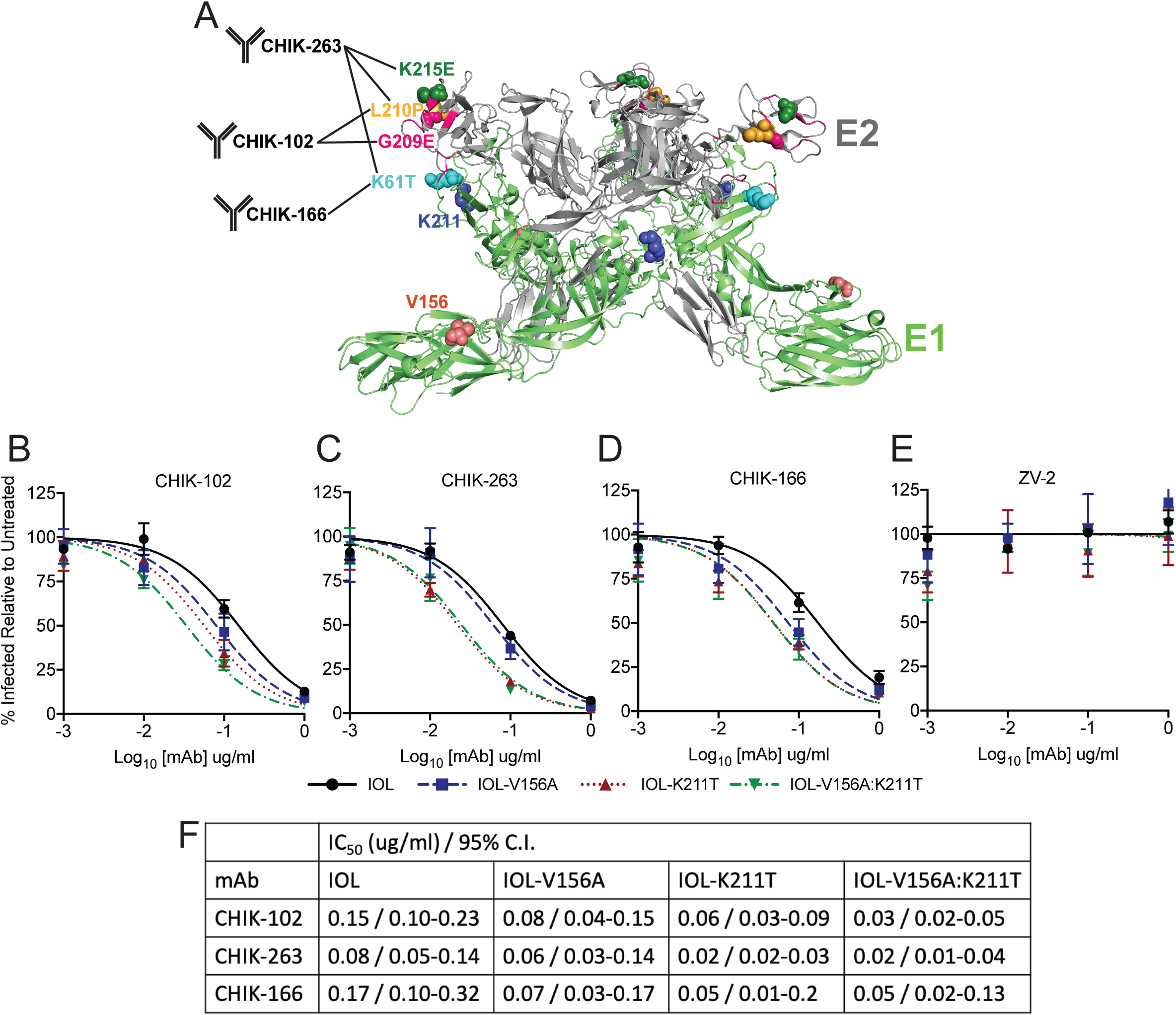
CHIKV E1 V156A and K211T impact antibody neutralization. (A) Anti-CHIKV mAbs target epitopes on E1 and E2 based on previously mapped escape mutants, labelled and colored spheres [42]. Pink ribbon represents CHIK-263 Fab footprint as previously mapped by Cryo-EM [43]. Virus neutralization assays using E2-targeting mAbs CHIK-102 (B) and CHIK-263 (C), E1-targeting mAb CHIK-166 (D), and ZIKV E-targeting control mAb ZV-2 (E) were conducted by incubating ZsGreen-expressing variants (MOI =1) with mAbs at indicated concentrations for 30 minutes and mixtures were used to infect BHK-21 cells. Cells were incubated for 18 hours, fixed, DAPI stained, and the percent of infected cells relative to untreated controls was quantified using fluorescence imaging. Data is representative of three independent experiments performed in duplicate. Non-linear regression was used to determine best fit curves, IC_50_ values and confidence intervals (F).

### CHIKV E1 K211T promotes dissemination in *Aedes aegypti* mosquitoes

Previous studies described the isolation of novel CHIKV isolates of the Asian genotype carrying the mutation K211E from regions of South East Asia [44–46]. Noting that these regions are *Ae. aegypti*-dominated, a subsequent study showed K211E increases fitness in *Ae. aegypti* [47]. To test whether K211T and/or V156A would be advantageous in *Ae. aegypti in vivo*, we infected *Ae. aegypti* mosquitoes with either of the single mutants or the double mutant in the IOL background and measured infectious titers in the bodies as well as disseminated virus in the legs and wings. IOL-V156A:K211T had slightly lower titers in the bodies of mosquitoes, suggesting decreased infectivity in *Ae. aegypti* (Figure 7A). However, IOL-K211T had elevated titers in the legs and wings, possibly corroborating previous findings that associated residue 211 with an advantage in *Ae. aegypti* (Figure 7B). The percentage of mosquitoes with infected bodies that also had disseminated virus in the legs and wings (% dissemination) was not statistically significant between the viruses (Figure 7C). Though the differences observed were modest, the varying phenotypes between V156A, K211T, and the double mutant in *Ae. aegypti* support the interplay of these E1 residues during CHIKV infection of mosquito vectors.

**Figure 7:**
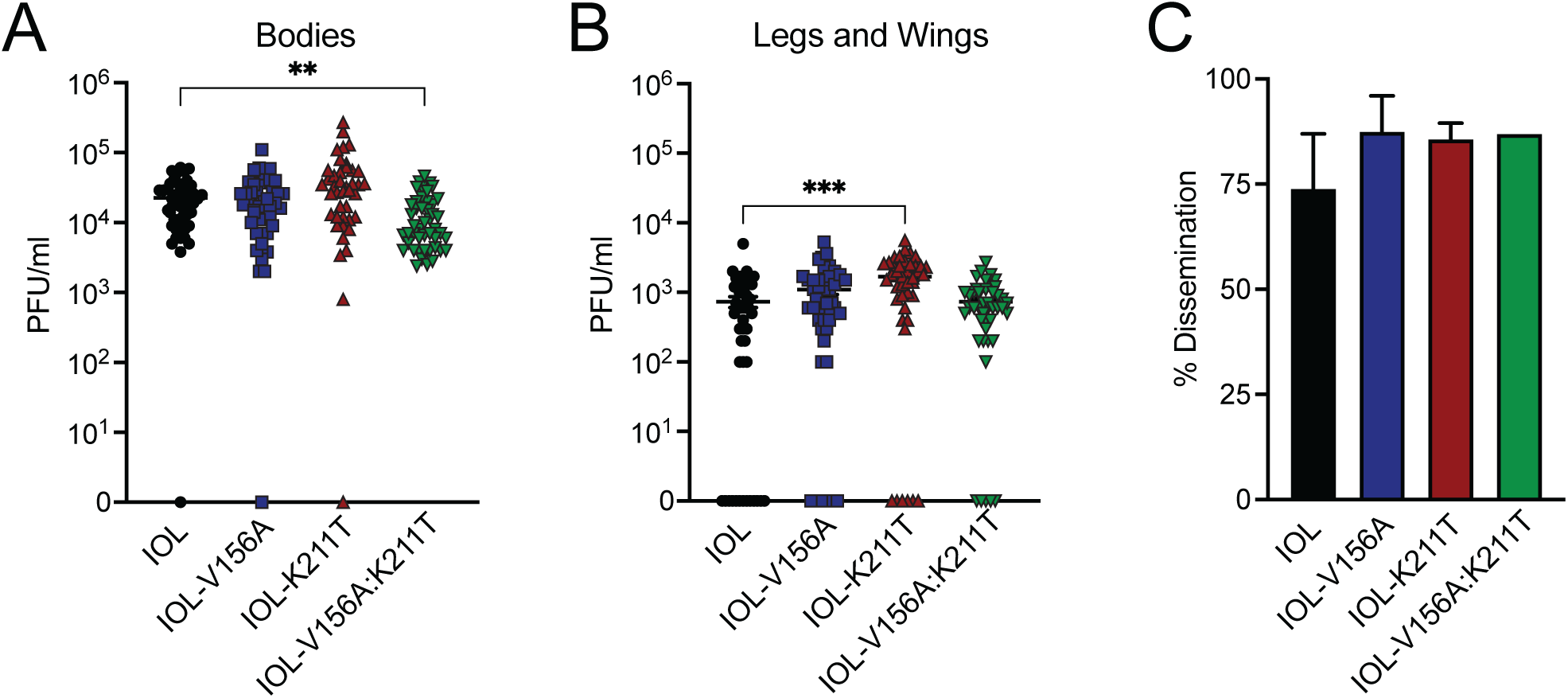
CHIKV K211E enhances dissemination in *Ae. aegypti* mosquitoes. Infectious titers quantified in bodies (A) and legs and wings (B) of *Ae. aegypti* mosquitoes infected with wild-type CHIKV or E1 variants (IOL V156A, N=45; IOL-K211T, N=42; IOL V156A+K211T, N=46). 7-day-old *Ae. aegypti* mosquitoes were fed an artificial blood meal containing virus (10^6^ PFU/ml) and incubated for 14 days. Viral titers were determined for homogenized mosquito bodies or legs/wings by plaque assay. Percent dissemination was determined as the percent of mosquitoes for which titers were detected in the body as well as legs and wings (C). Data represent two independent experiments, * p < 0.05, ** p < 0.01, *** p < 0.001; Kruskal-Wallis test.

### CHIKV E1 V156A and K211T enhance viral replication and pathogenesis in mice

Finally, we sought to determine the impact of V156A and K211T during infection in a mouse model. We infected 4 to 7-week-old C57BL/6 mice subcutaneously via the footpad with each of the CHIKV variants on the IOL background, as it was a variant on this background that exhibited an entry advantage *in vitro*. At 2 days post infection, during peak viremia [48], we measured foot swelling and quantified infectious titers in serum and in the injected foot. While each variant reached similar levels in serum, all three variants had higher titers than wild-type CHIKV in the injected foot (Figure 8A). Infection with all three variants also resulted in increased swelling of the foot, with the double mutant having the greatest impact on swelling (Figure 8B). This demonstrates that discrete E1 residues can impact both viral replication and virus-induced pathology in the host.

**Figure 8:**
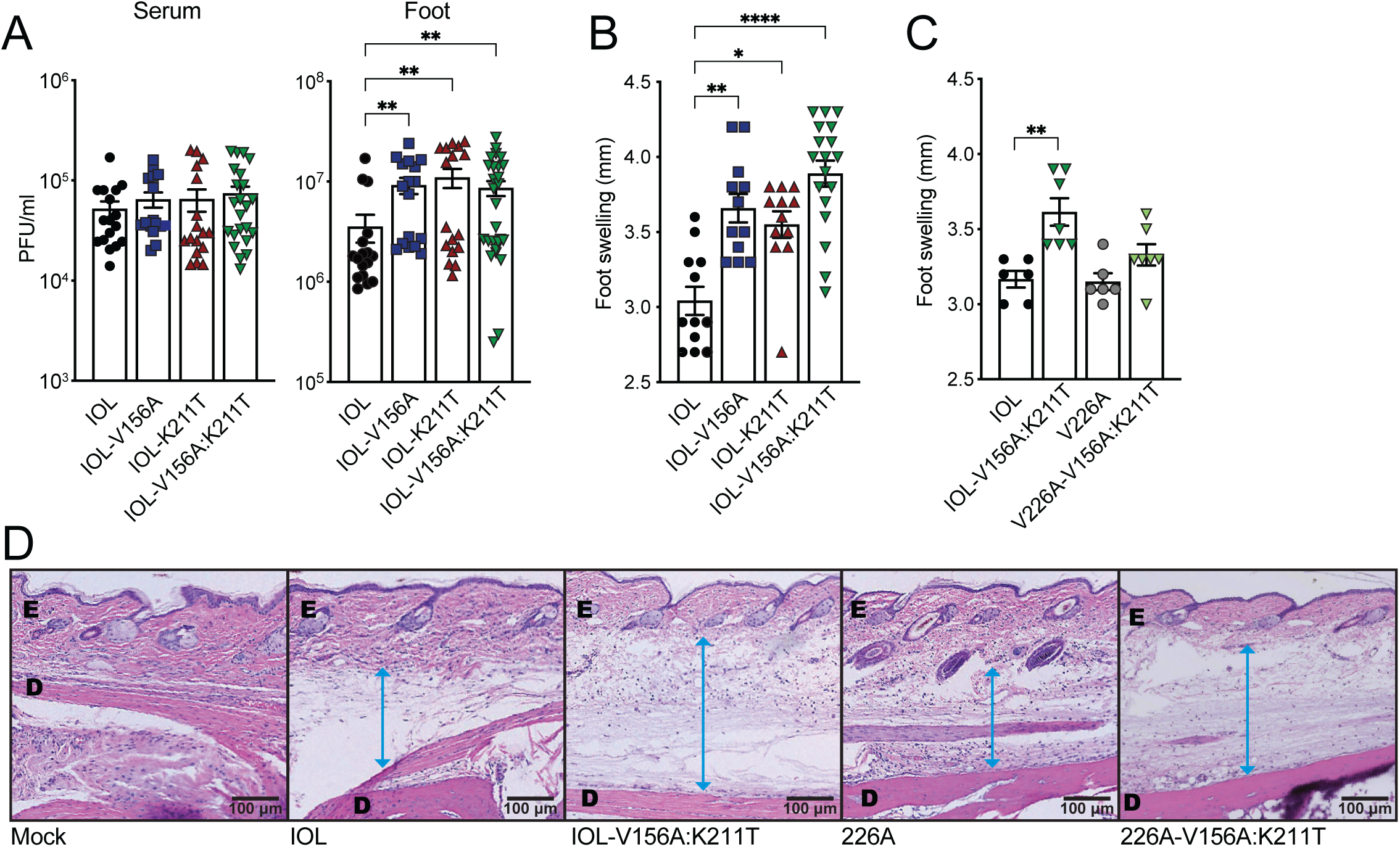
CHIKV V156A and K211T increase replication and inflammation in mice. 4-7-week-old C57BL/6 mice were infected via subcutaneous injection of the footpad with 1000 PFU of virus. Quantification of infectious titers in the serum and ipsilateral foot (A) and foot swelling (B and C) at 2 dpi for mice infected with wild-type CHIKV or E1 variants (IOL V156A, N=7-17; IOL-K211T, N=12-17; IOL V156A+K211T, N=7-27).. Pathology of the feet was visualized by H&E staining (D). ‘E’= epidermis; ‘D’= dermis. Blue arrow indicates the measurement of swelling of the area between the dermis and epidermis. Data represent at least two independent experiments, * p < 0.05, ** p < 0.01, *** p < 0.001; Kruskal-Wallis test.

Considering that we observed differing phenotypes in *in vitro* assays whether V156A and K211T were expressed on the 226V or 226A-expressing backgrounds, we questioned whether residue 226 would also contribute to the increased inflammation observed in mice. To address this, we infected mice with the V156A and K211T double mutant on either the IOL or V226A background and measured foot swelling at 2 days post infection. The two parental viruses caused similar swelling whereas V226A-V156A:K211T caused intermediate swelling compared with the parental viruses and IOL-V156A:K211T (Figure 8C). Histological examination of the ipsilateral foot showed an increase in subcutaneous swelling in mice infected with V156A and K211T viruses, which was more pronounced for the double mutant in the IOL background, as was suggested by the swelling measurements (Figure 8D), indicating a functional link between E1 residues 156, 211 and 226 that impacts replication and pathology in a mammalian host.

## DISCUSSION

Alphaviruses are prominent disease-causing arboviruses. They will remain an ongoing issue as long as the vectors that transmit them exist around human populations and continue to expand in geographical abundance [49]. This raises our need to understand the mechanistic details of how these viruses are transmitted and cause disease. Using an *in vivo* transmission system in the lab, we identified a frequently occurring mutation in the CHIKV E1 glycoprotein, V156A, that subsequently emerged in nature, co-occurring with the E1 mutation K211T. In this work, we characterize these mutations *in vitro* and *in vivo* and show that these residues function in virus binding and entry and can significantly impact dissemination in mosquitoes and pathology in mice.

We demonstrated that the E1 mutations V156A, K211T, and the double mutant exhibit decreased cell binding and decreased *in vitro* binding to heparin. Interestingly, these effects were observed to a lesser extent in the background of V226A. Residue 226 has previously been associated with host-specific and epistatic functions and our study further implicates 226 in modulating phenotypes driven by other E1 residues [28, 31, 47]. These results are intriguing as CHIKV is known to use glycosaminoglycans as attachment factors, yet these interactions, so far, have been attributed to residues of the E2 glycoprotein [14, 50, 51]. The CHIKV strain 181/25, for instance, displays increased GAG binding attributed to the E2 mutation G82R, which was acquired following passage in cell culture [11, 14, 50]. Importantly, natural isolates of CHIKV are dependent on GAGs for efficient infection of some mammalian cells. Recent analysis of glycan interactions for diverse CHIKV strains revealed members of each genetic clade preferentially bind GAGs over other glycans [14, 16, 52]. However, strains from different clades displayed different binding to GAGs and this variation indicates the potential array of genetic differences that may drive GAG-virus interactions.

One potential explanation for reduced attachment and GAG interactions may be through changes in the glycoprotein complex structure. Using monoclonal antibodies, we observed that E1 V156A and K211T can alter antibody neutralization. These results suggest that there are structural changes to the mature virion caused by residue changes in the E1 glycoprotein. Within the glycoprotein lattice of the mature particle, E2 makes only intra-spike contacts between E1-E2 whereas biochemical and structural studies have identified the inter-glycoprotein spike interface to consist of contacts between neighboring spikes of the mature particle to be sustained exclusively by E1, forming the icosahedral scaffolding and joining the 80 glycoprotein spikes [53–56]. The E1 residues 156 and 211 are located within this E1-E1 interface between adjacent E1-E2 trimeric spikes (Figure 9A, squares). These E1 mediated interactions being important for stabilization of the glycoprotein lattice and for glycoprotein spike orientation together with E1-E2 intra-spike contacts make E1 well poised to influence how the particle engages with the cell surface. In the way that E1 V156A and K211T have a far-reaching impact on binding by mAbs that target outward facing residues of E2, they could similarly have an impact on the orientation of regions known to engage GAGs, such as that surrounding E2 position 82, located in the upward-facing inner cavity of the spike (Figure 9B). Inspecting the electrostatic potential of the inter-spike interface, it is notable that the interface harbors many exposed positively charged patches that could potentially favor GAG-protein interactions that are mostly ionic (Figure 9C) [57].

**Figure 9:**
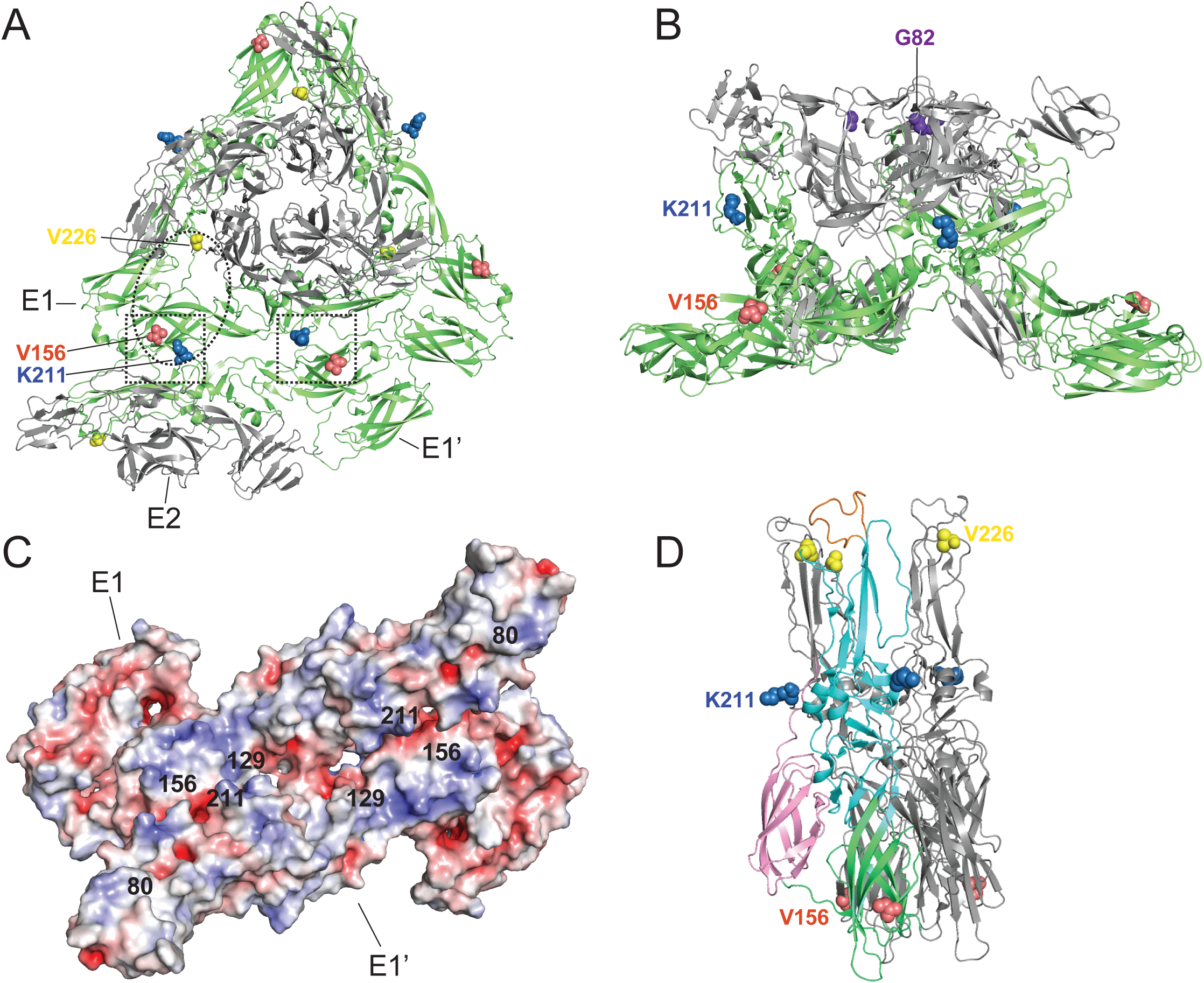
Structural visualization of E1 residues 156 and 211. (A) Top view of CHIKV E1/E2 spike and E1-E1 inter-spike interface (PDB 2XFB). Squares denote positions 156 and 211 that face each other on E1 and E1’. Oval denotes approximate region predicted to interact with Mxra8 [18] (B) Side view of E1/E2 spike (PDB 2FXB). (C) Electrostatic potentials of E1-E1 inter-spike interface. E1 residues of interest are indicated. Negative electrostatic potential is shown in red and positive electrostatic potential in blue. (D) Semliki Forest Virus E1-E1 Post-fusion spike (PDB 1RER) with one E1 monomer colored by domain (green= domain I, cyan= domain II, pink = domain III). The E1 fusion loop is shown in orange.

In addition to interactions with GAGs, E1 residues 156, 211, and 226 each lie within the predicted contact zones of Mxra8 (Figure 9A, oval) [13, 18]. Notably, the recent characterization of CHIKV-GAG interactions found Mxra8 expression to be inversely correlated to GAG-dependency for CHIKV binding and infection [16]. Also intriguing, considering our data, is that *in vivo*, Mxra8 deficient mice exhibit a modest decrease in titers at the sight of injection following infection via footpad, but dramatically decreased foot swelling [17]. In our study, we find V156A and K211T to increase foot swelling and that this is regulated by residue 226. Taken together, E1 residues in the inter-spike interface may play critical roles in glycoprotein assembly, spike orientation, and host-pathogen interactions at the cell membrane. It will be insightful to investigate the impact of E1 variants on Mxra8 interactions, in addition to GAG usage, in future studies.

Residue 226 has also been shown to modulate CHIKV fusion and cholesterol dependence and to possess a functional link to residues near the fusion loop [28, 58]. Visualizing E1 residues 156 and 211 on the structurally similar Semliki Forest virus E1 protein that has been crystalized in its post-fusion trimer formation shows 156 is in the dynamic E1 hinge and near the domain I-domain III linker, a flexible region important for refolding, trimerization, and successful fusion [36] (Figure 9D). Position 211 is in a contact region between E1 monomers within the trimer [59]. This region is outward facing and possibly critical for the cooperative ring formation that occurs between up to 6 interacting E1 trimers during fusion [24, 25]. In this study, IOL-V156A:K211T consistently infected more cells at low pH in a fusion assay compared with wild-type virus, which was not observed with V226A-V156A:K211T. V156A emerged in both V226 and A226 backgrounds, but only co-occurred with K211T in the A226 background. Neither position 156 or 211 are located near the fusion loop and it is intriguing that they further demonstrate a long-range functional link to residue 226.

Finally, we found E1 V156A and K211T to impact virulence in adult C57BL/6 mice. Elevated titers and swelling of the inoculated foot were observed for V156A and K211T variants and as we hypothesized, this was dictated by residue 226, as variant V226A-V156A:K211T exhibited intermediate swelling. Previous work has shown that mutations at E2 residue 82, which impacts GAG utilization, also modulates virus infection and arthritis in mice [50]. Therefore, our findings are in line with these studies and may suggest that modulating GAG interactions through E1 or E2 can impact inflammation in mice. Future studies will investigate the underlying mechanism of this phenotype and potential influences on the immune response, including possible differential recruitment of cellular infiltrates and expression of inflammatory cytokines. Multiple recently identified E1 mutations found to increase pathogenesis, are located in the E1-E1 inter-spike interface where residue 156 and 211 are located, potentially highlighting an evolution hotspot for emerging CHIKV variants (Figure 9C).

This work has provided evidence for functional roles of emerging CHIKV E1 variants V156A and K211T, with implications in cell attachment and pathogenesis. We demonstrate that the function of E1 in cell entry extends beyond membrane fusion and that discrete E1 regions can also influence cell binding. As suggested by altered neutralization by mAbs, this influence is potentially mediated by structural changes to the particle. Our findings also highlight V156A and K211T as determinants of CHIKV virulence with evidence that E1 can regulate swelling at the site of infection in the host, a dynamic that will be important to further elucidate. Altogether, these results expand our current understanding of the multi-functionality of the E1 glycoprotein, which will be useful for the development of therapeutic and preventative tools.

## MATERIALS AND METHODS

### Cell lines

Mammalian cell lines were maintained at 37°C in 5% CO_2_. Baby hamster kidney (BHK-21, ATCC CCL-10) were grown in Dulbecco’s modified Eagle’s medium (DMEM; Corning) supplemented with 10% fetal bovine serum (FBS, Atlanta Biologicals) and 1% non-essential amino acids (NEAA, Gibco). Vero cells (ATCC CCL-81) were grown in DMEM supplemented with 10% newborn calf serum (NBCS, Gibco) and 1% penicillin/streptomycin (Corning). Mosquito cell lines were maintained at 28°C in 5% CO_2_. *Ae. aegypti* cells (Aag2, provided by P. Turner, Yale University) were grown in DMEM supplemented with 10% FBS and 1% NEAA. *Ae. albopictus* cells (C6/36, ATCC CRL-1660) were maintained in L-15 medium (Corning) supplemented with 10% FBS, 1% tryptose phosphate broth (Invitrogen), and 1% NEAA.

### Viruses

Wild-type chikungunya virus (CHIKV) and E1 glycoprotein variants were generated from the CHIKV strain 06-049 (AM258994) infectious clone, described previously [60]. Amino acid substitutions were introduced into the E1 glycoprotein by site-directed mutagenesis using a cloning plasmid that contained the genomic region of interest flanked by XhoI and NotI restriction enzyme sites, Phusion DNA polymerase (Thermo-Fisher) and the primers in Table 1. The XhoI/NotI fragment was then sub-cloned into the full-length infectious clone plasmid using the same restriction sites. The variants were also introduced by this method to a ZsGreen-expressing CHIKV infectious clone described previously [28]. All E1 variants were confirmed by full genome Sanger sequencing. To produce *in vitro* transcribed viral RNA, 10 μg of each CHIKV plasmid was linearized overnight using NotI (Invitrogen), purified by phenol:chloroform extraction and used as a template for *in vitro* transcription using the SP6 mMessage mMachine kit (Invitrogen). Resulting RNA was purified by phenol:chloroform extraction, aliquoted and stored at -80°C. To produce stocks of infectious virus, 3.9 x 10^6^ BHK-21 cells were mixed with 10 μg RNA, electroporated with one pulse at 1200 V, 25 μF and infinite resistance then incubated at 37°C for 72 hours. Virus-containing supernatant, passage 0 (P0), was collected, clarified by centrifugation for 5 minutes at 1,200 RPM and used to infect a monolayer of BHK-21 cells for 24 hours to produce passage 1 (P1). P1 supernatant, used as the working virus stock, was collected, clarified by centrifugation for 5 minutes at 1,200 RPM, aliquoted and stored at - 80°C. Gradient-purified virus stocks were produced as described above, with the addition of ultracentrifugation of P1 over a 20% sucrose cushion at 25,000 RPM for 2 hours and resuspended in infection media (DMEM, 0.2% BSA and 1 mM HEPES pH 7.4,) before aliquoting and storing at -80°C. Viral titers were determined by plaque assays on Vero cells as described below. Full genomes of all P1 stocks were Sanger sequenced (Genewiz) to ensure the absence of second-site mutations. All CHIKV infections were conducted at the NYU Grossman School of Medicine under biosafety level 3 conditions.

**Table 1.**
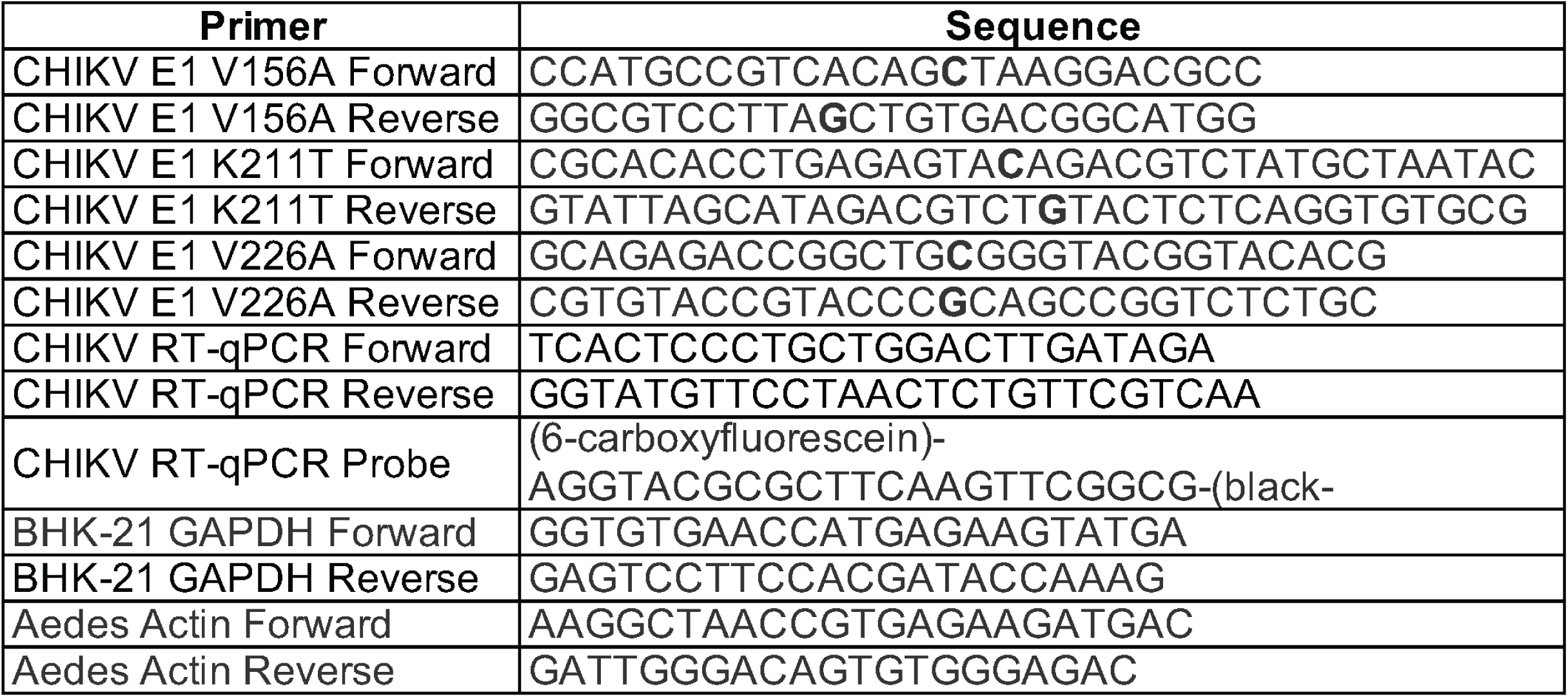
PCR primers used in this study.

### Plaque assay

400,000 Vero cells/well were seeded in 12-well plates one day prior to infection. Ten-fold dilutions of virus-containing samples were prepared in DMEM and 200 μl of each dilution was used to infect cell monolayers. Virus-cell mixtures were incubated for 1 hour at 37°C then overlaid with DMEM containing 2% NBCS and 0.8% agarose. Cells were incubated for 3 days at 37°C then fixed with 4% formalin and plaques were visualized using crystal violet staining. Viral titers were determined by counting the number of plaques on the lowest countable dilution.

### CHIKV RNA extraction and genome quantification

RNA was purified using TRIzol (Fisher-Scientific) following the manufacturer’s instructions. CHIKV genomes/mL were quantified by RT-qPCR using the Applied Biosystems TaqMan RNA-to-CT One-Step Kit (Fisher-Scientific) with primers in Table 1. A CHIKV RNA standard was generated from *in vitro* transcribed RNA as described above and used to calculate CHIKV genomes/mL. All RT-qPCR analyses were run with a CHIKV standard and all samples and standards were run in technical duplicate.

### Fusion-from-without assay

BHK-21 cells were incubated at 4°C in binding buffer (RPMI, 0.2% BSA, 10 mM HEPES, 20 mM NH_4_Cl) for 1 hour. Gradient-purified ZsGreen-expressing viruses were diluted in binding buffer and allowed to bind the cell monolayer at 4°C for 1 hour at an MOI of 1. Unbound virus was removed and fusion was induced at a range of pHs by adding fusion buffer (RPMI, 0.2% BSA, 10 mM HEPES, 30 mM succinic acid) adjusted to each pH. After 2 minutes, fusion buffer was removed and complete media supplemented with 20 mM NH_4_Cl was added and cells were incubated at 37°C for 16 hours. Cells were then fixed with 2% PFA and DAPI (Thermo-Scientific) stained. Infected cells were quantified using a CellInsight CX7 high-content microscope (Thermo-Scientific) and HCS Navigator Software Version 6.6.1 (Thermo-Scientific).

### Lysosomotropic agent sensitivity assay

BHK-21 cells were incubated in DMEM containing a range of concentrations of NH_4_Cl or Bafilomycin A1 for 3 hours. Cells were then infected at an MOI of 1 for 1 hour with ZsGreen-expressing viruses in the presence of either lysosomotropic agent. Virus was removed and cells were washed with phosphate buffer saline (PBS) three times and replenished with complete media supplemented with either agent. Cells were incubated at 37°C for 16 hours then fixed, and DAPI stained. Infected cells were quantified using fluorescent microscopy as described above.

### Virus-cell binding assay

BHK-21, Aag2, or C6/36 cells were incubated in binding buffer at 4°C for 1 hour. Purified viruses were diluted in binding buffer to an infection ratio of 10 genomes/cell and allowed to bind cells at 4°C for one hour. Unbound virus was removed and cells were washed with PBS three times. Cells were harvested in TRIzol and total RNA was purified following the manufacturer’s instructions. cDNA was synthesized using the Maxima H RT Kit (Invitrogen) and viral genomes relative to cellular GAPDH (BHK-21) or actin (C6/36 and Aag2 cells) were quantified using SYBR Green qPCR with primers targeting CHIKV nsP4, listed in Table 1.

### Heparin ELISA

96-well ELISA plates coated in biotin-conjugated heparin were incubated with serial diluted gradient-purified virus particles (10^8^-10^5^ genomes) for 1-2 hours at room temperature. Unbound virus was removed and plates were washed three times. Bound virus was detected by incubation with a primary anti-CHIKV E2 antibody and an HRP-conjugated secondary antibody, followed by addition of TMB substrate for up to 15 minutes. Oxidation of TMB was stopped by adding sulfuric acid and absorbance at 450 nm was measured using a plate reader. Heparin ELISAs were performed at the University of Pittsburgh under biosafety level 3 conditions.

### Antibody neutralization assay

ZsGreen-expressing viruses were incubated for 30 minutes at room temperature with a range of concentrations of monoclonal antibodies targeting CHIKV E1 (CHIK-166), E2 (CHIK-102 and CHIK-263) [42] or Zika Virus E as a control (ZV-2, Sigma-Aldrich) (kindly provided by Dr. Michael Diamond, Washington University). Virus and antibody mixtures were used to inoculate BHK-21 cells, which were then incubated at 37°C for 18 hours. Cells were fixed, DAPI-stained and infected cells were quantified as described above.

### Mouse infections

4-7-week-old male and female C57BL/6 mice were infected in the left rear foot pad with 1000 PFU of wild-type CHIKV and each variant diluted in 50 μl of PBS. At 2 days post infection, foot swelling was measured using calipers and mice were euthanized. Blood was collected by cardiac puncture and harvested organs were homogenized in 500 μl PBS with two 5 mm stainless steel beads using a tissue lyser (Tissue-Lyser II, Qiagen) for two 2-minute rounds at 30 Hz and centrifuged to pellet debris for 10 minutes at 8,000 rpm. Infectious particles in the supernatants were quantified by plaque assay as described above. Animal experiments were performed under biosafety level 3 conditions in accordance with all NYU School of Medicine Institutional Animal Care and Use Committee guidelines (IACUC) (Protocol # IA16-01783).

### Mosquito infections

*Ae. aegypti* mosquitoes (Poza Rica, Mexico, P20) were obtained from Dr. Gregory Ebel (Colorado State University) [61]. Mosquitoes were reared and maintained in Memmert humidified chambers at 28°C and 70% humidity with a 12-hour diurnal light cycle. Artificial blood meals were prepared by diluting viruses to 10^6^ PFU/mL in washed sheep whole blood (Fisher-Scientific) supplemented with 5 mM ATP and fed to 7-day old female mosquitoes for 60 minutes through a pork intestine membrane warmed to 37°C. Engorged females were sorted and incubated for 14 days in 28°C chambers while fed 10% sucrose *ad libitum*. Mosquito bodies and legs/wings were removed, placed in 250 μL PBS containing a 5 mm stainless steel bead and homogenized using a tissue lyser (Tissue-Lyser II, Qiagen) for 2-minutes at 30 Hz. Debris was pelleted by centrifugation at 8,000 RPM for 10 minutes and infectious titers of the supernatants were determined by plaque assay as described above.

### Western blotting

10^5^ PFU of sucrose-gradient purified virions were suspended in Laemmli sample buffer supplemented with 2-mercaptoethanol. Suspensions were boiled at 95°C for 10 minutes and centrifuged at 10,000 x g for 1 minute. Protein was separated by SDS-PAGE and transferred to a polyvinylidene difluoride (PVDF) membrane (Immobilon, Millipore). Blots were blocked using 5% milk in Tris-buffered saline containing 0.1% Tween 20 (TBS-T). Blots were incubated with primary antibodies to CHIKV E1 (provided by Dr. Gorben Pijlman), CHIKV E2 (CHIK-187 provided by Dr. Michael Diamond), and CHIKV capsid (CHIK-122 provided by Dr. Andres Merits). Blots were then washed extensively and incubated with HRP-conjugated goat anti-mouse or anti-rabbit IgG secondary antibodies (Invitrogen). Blots were developed using the SuperSignal West Pico PLUS chemiluminescent substrate kit (Thermo) and imaged using the ChemiDoc MP Imaging System (BioRad).

### Electron microscopy

Purified viruses were fixed in 4% paraformaldehyde in PBS overnight at 4°C. 5 μl of fixed viruses were added onto glow discharged carbon coated 400 mesh Cu/Rh grid (Ted Pella Inc., Redding, CA), and stained with 1% aqueous uranyl acetate (Polysciences, Inc, Warrington, PA). Stained grids were imaged under Talos120C transmission electron microscope (Thermo Fisher Scientific, Hillsboro, OR) using Gatan OneView digital camera (4K x 4K, Gatan, Inc., Pleasanton, CA).

### Histopathology

The method used for preparation of samples for histology was adapted from a protocol kindly provided by Dr. Deborah Lenschow (Washington University). At 2 days post infection, the injected left rear feet of mice were harvested and fixed in 10% neutral buffered formalin for 72 hours. Feet were rinsed with PBS and decalcified using 5 M EDTA for 2 weeks at 4°C. Feet were rinsed and stored in 70% ethanol. Feet were embedded in paraffin and 5 μm sections were prepared at the NYU School of Medicine Experimental Pathology Research Laboratory. Sections were stained with hematoxylin and eosin (H&E) and imaged under light microscopy.

### Protein structures

Protein structural data was accessed via Protein Data Bank (PDB) and analyzed using PyMOL version 2.3.3. Surface electrostatic potential maps were generated using PyMOL plug-in Adaptive Poisson-Boltzmann Solver (APBS) [62].

### Statistics and data analysis

All data and statistical analyses were performed using GraphPad Prism software (version 9.0.0). Two-way ANOVA with Bonferroni’s multiple comparisons test, Kruskal-Wallis test with Dunn’s multiple comparisons test, Mann-Whitney U test, and non-linear regression were performed as indicated in the figure legends. All experiments were performed at least two independent times with internal duplicates. P<0.05 is considered significant.

## Acknowledgements

We thank all members of the Stapleford Lab for valuable discussions, Dr. Meike Dittmann at the NYU Grossman School of Medicine for use of the CX7 high-content microscope, the Office of Science & Research High-Containment Laboratories at NYU Grossman School of Medicine for their support in the completion of this research, and Dr. Alice Liang and the NYU Langone Health DART Microscopy Lab for consultation and assistance with TEM work. This work was supported by a start-up package from the NYU Grossman School of Medicine (K.A.S), the Public Health Service Institutional Research Training Award T32 AI007180 (M.V.R), the American Heart Association Postdoctoral Fellowship (19-A0-00-1003686) (M.G.N), NYU Cancer Center Support Grants NIH/NCI P30CA016087 (Microscopy Lab) and NIH/NCI 5 P30CA16087 (Experimental Pathology Research Laboratory), F31 AI147440 (N.M.), and start-up funds provided by the Department of Pediatrics at the University of Pittsburgh (L.A.S).

## References

1. Weaver, S.C., et al., Zika, Chikungunya, and Other Emerging Vector-Borne Viral Diseases. Annu Rev Med, 2018. 69: p. 395–408.

2. Yactayo, S., et al., Epidemiology of Chikungunya in the Americas. J Infect Dis, 2016. 214(suppl 5): p. S441–S445.

3. Gill, C.M., et al., Five Emerging Neuroinvasive Arboviral Diseases: Cache Valley, Eastern Equine Encephalitis, Jamestown Canyon, Powassan, and Usutu. Semin Neurol, 2019. 39(4): p. 419–427.

4. Levi, L.I. and M. Vignuzzi, Arthritogenic Alphaviruses: A Worldwide Emerging Threat? Microorganisms, 2019. 7(5).

5. Adouchief, S., et al., Sindbis virus as a human pathogen-epidemiology, clinical picture and pathogenesis. Rev Med Virol, 2016. 26(4): p. 221–41.

6. Weaver, S.C. and N.L. Forrester, Chikungunya: Evolutionary history and recent epidemic spread. Antiviral Res, 2015. 120: p. 32–9.

7. Control, E.C.f.D.P.a. Geographical distribution of chikungunya virus disease cases reported worldwide, 2020. 2021 June 1, 2021]; Available from: https://www.ecdc.europa.eu/en/publications-data/geographical-distribution-chikungunya-virus-disease-cases-reported-worldwide-2020.

8. Pietilä, M.K., K. Hellström, and T. Ahola, Alphavirus polymerase and RNA replication. Virus Research, 2017. 234: p. 44–57.

9. Jose, J., J.E. Snyder, and R.J. Kuhn, A structural and functional perspective of alphavirus replication and assembly. Future Microbiol, 2009. 4(7): p. 837–56.

10. Gardner, C.L., et al., Deliberate attenuation of chikungunya virus by adaptation to heparan sulfate-dependent infectivity: a model for rational arboviral vaccine design. PLoS neglected tropical diseases, 2014. 8(2): p. e2719.

11. Levitt, N.H., et al., Development of an attenuated strain of chikungunya virus for use in vaccine production. Vaccine, 1986. 4(3): p. 157–162.

12. Prado Acosta, M., et al., Surface (S) layer proteins of Lactobacillus acidophilus block virus infection via DC-SIGN interaction. Frontiers in microbiology, 2019. 10: p. 810.

13. Zhang, R., et al., Mxra8 is a receptor for multiple arthritogenic alphaviruses. Nature, 2018: p. 1.

14. Silva, L.A., et al., A single-amino-acid polymorphism in Chikungunya virus E2 glycoprotein influences glycosaminoglycan utilization. Journal of virology, 2014. 88(5): p. 2385–2397.

15. Klimstra, W.B., K.D. Ryman, and R.E. Johnston, Adaptation of Sindbis virus to BHK cells selects for use of heparan sulfate as an attachment receptor. J Virol, 1998. 72(9): p. 7357–66.

16. McAllister, N., et al., Chikungunya virus strains from each genetic clade bind sulfated glycosaminoglycans as attachment factors. Journal of virology, 2020. 94(24).

17. Zhang, R., et al., Expression of the Mxra8 Receptor Promotes Alphavirus Infection and Pathogenesis in Mice and Drosophila. Cell reports, 2019. 28(10): p. 2647–2658.e5.

18. Basore, K., et al., Cryo-EM structure of Chikungunya virus in complex with the Mxra8 receptor. Cell, 2019. 177(7): p. 1725–1737. e16.

19. Bernard, E., et al., Endocytosis of chikungunya virus into mammalian cells: role of clathrin and early endosomal compartments. PloS one, 2010. 5(7): p. e11479–e11479.

20. Hoornweg, T.E., et al., Dynamics of chikungunya virus cell entry unraveled by single- virus tracking in living cells. Journal of virology, 2016. 90(9): p. 4745–4756.

21. Wahlberg, J., W. Boere, and H. Garoff, The heterodimeric association between the membrane proteins of Semliki Forest virus changes its sensitivity to low pH during virus maturation. Journal of virology, 1989. 63(12): p. 4991–4997.

22. Wahlberg, J.M. and H. Garoff, Membrane fusion process of Semliki Forest virus. I: Low pH-induced rearrangement in spike protein quaternary structure precedes virus penetration into cells. The Journal of cell biology, 1992. 116(2): p. 339–348.

23. Wahlberg, J.M., et al., Membrane fusion of Semliki Forest virus involves homotrimers of the fusion protein. Journal of Virology, 1992. 66(12): p. 7309–7318.

24. Gibbons, D.L., et al., Conformational change and protein–protein interactions of the fusion protein of Semliki Forest virus. Nature, 2004. 427(6972): p. 320.

25. Gibbons, D.L., et al., Visualization of the Target-Membrane-Inserted Fusion Protein of Semliki Forest Virus by Combined Electron Microscopy and Crystallography. Cell, 2003. 114(5): p. 573–583.

26. Byrd, E.A. and M. Kielian, The Alphavirus E2 Membrane-Proximal Domain Impacts Capsid Interaction and Glycoprotein Lattice Formation. Journal of Virology, 2018: p. JVI.01881-18.

27. Quiroz, J.A., et al., Human monoclonal antibodies against chikungunya virus target multiple distinct epitopes in the E1 and E2 glycoproteins. PLoS pathogens, 2019. 15(11): p. e1008061.

28. Noval, M.G., et al., Evolution-Driven Attenuation of Alphaviruses Highlights Key Glycoprotein Determinants Regulating Viral Infectivity and Dissemination. Cell Rep, 2019. 28(2): p. 460–471 e5.

29. Coffey, L.L., et al., Factors shaping the adaptive landscape for arboviruses: implications for the emergence of disease. Future Microbiol, 2013. 8(2): p. 155–76.

30. Anishchenko, M., et al., Venezuelan encephalitis emergence mediated by a phylogenetically predicted viral mutation. Proceedings of the National Academy of Sciences, 2006. 103(13): p. 4994–4999.

31. Tsetsarkin, K.A., et al., A single mutation in chikungunya virus affects vector specificity and epidemic potential. PLoS pathogens, 2007. 3(12): p. e201.

32. Sam, I.-C., et al., Chikungunya virus of Asian and central/east African genotypes in Malaysia. Journal of Clinical Virology, 2009. 46(2): p. 180–183.

33. Tsetsarkin, K.A., et al., Chikungunya virus emergence is constrained in Asia by lineage- specific adaptive landscapes. Proceedings of the National Academy of Sciences, 2011: p. 201018344.

34. Souza, T.M., et al., First Report of the East-Central South African Genotype of Chikungunya Virus in Rio de Janeiro, Brazil. PLoS Curr, 2017. 9.

35. Stapleford, K.A., et al., Emergence and transmission of arbovirus evolutionary intermediates with epidemic potential. Cell host & microbe, 2014. 15(6): p. 706–716.

36. Zheng, Y., et al., The domain I-domain III linker plays an important role in the fusogenic conformational change of the alphavirus membrane fusion protein. Journal of virology, 2011: p. JVI. 00596-11.

37. Glomb-Reinmund, S. and M. Kielian, The role of low pH and disulfide shuffling in the entry and fusion of Semliki Forest virus and Sindbis virus. Virology, 1998. 248(2): p. 372–81.

38. Helenius, A., M. Marsh, and J. White, Inhibition of Semliki Forest Virus Penetration by Lysosomotropic Weak Bases. Journal of General Virology, 1982. 58(1): p. 47–61.

39. Van Deurs, B., P.K. Holm, and K. Sandvig, Inhibition of the vacuolar H (+)-ATPase with bafilomycin reduces delivery of internalized molecules from mature multivesicular endosomes to lysosomes in HEp-2 cells. European journal of cell biology, 1996. 69(4): p. 343–350.

40. Klimstra, W.B., K.D. Ryman, and R.E. Johnston, Adaptation of Sindbis virus to BHK cells selects for use of heparan sulfate as an attachment receptor. Journal of virology, 1998. 72(9): p. 7357–7366.

41. Bernard, K.A., W.B. Klimstra, and R.E. Johnston, Mutations in the E2 glycoprotein of Venezuelan equine encephalitis virus confer heparan sulfate interaction, low morbidity, and rapid clearance from blood of mice. Virology, 2000. 276(1): p. 93–103.

42. Pal, P., et al., Development of a highly protective combination monoclonal antibody therapy against Chikungunya virus. PLoS pathogens, 2013. 9(4): p. e1003312.

43. Zhou, Q.F., et al., Structural basis of Chikungunya virus inhibition by monoclonal antibodies. Proceedings of the National Academy of Sciences of the United States of America, 2020. 117(44): p. 27637–27645.

44. Shrinet, J., et al., Genetic characterization of Chikungunya virus from New Delhi reveal emergence of a new molecular signature in Indian isolates. Virology Journal, 2012. 9(1): p. 100.

45. Sumathy, K. and K.M. Ella, Genetic diversity of chikungunya virus, India 2006–2010: Evolutionary dynamics and serotype analyses. Journal of Medical Virology, 2012. 84(3): p. 462–470.

46. Taraphdar, D. and S. Chatterjee, Molecular characterization of chikungunya virus circulating in urban and rural areas of West Bengal, India after its re-emergence in 2006. Transactions of The Royal Society of Tropical Medicine and Hygiene, 2015. 109(3): p. 197–202.

47. Agarwal, A., et al., Two novel epistatic mutations (E1:K211E and E2:V264A) in structural proteins of Chikungunya virus enhance fitness in Aedes aegypti. Virology, 2016. 497: p. 59–68.

48. Gardner, J., et al., Chikungunya virus arthritis in adult wild-type mice. Journal of virology, 2010. 84(16): p. 8021–8032.

49. Kraemer, M.U., et al., The global distribution of the arbovirus vectors Aedes aegypti and Ae. albopictus. elife, 2015. 4: p. e08347.

50. Ashbrook, A.W., et al., Residue 82 of the Chikungunya virus E2 attachment protein modulates viral dissemination and arthritis in mice. J Virol, 2014. 88(21): p. 12180–92.

51. Weber, C., et al., Identification of Functional Determinants in the Chikungunya Virus E2 Protein. PLoS neglected tropical diseases, 2017. 11(1): p. e0005318–e0005318.

52. Tanaka, A., et al., Genome-wide screening uncovers the significance of N-sulfation of heparan sulfate as a host cell factor for chikungunya virus infection. Journal of virology, 2017. 91(13): p. e00432–17.

53. Anthony, R.P. and D.T. Brown, Protein-protein interactions in an alphavirus membrane. Journal of Virology, 1991. 65(3): p. 1187–1194.

54. Ekström, M., P. Liljeström, and H. Garoff, Membrane protein lateral interactions control Semliki Forest virus budding. The EMBO Journal, 1994. 13(5): p. 1058–1064.

55. Pletnev, S.V., et al., Locations of carbohydrate sites on alphavirus glycoproteins show that E1 forms an icosahedral scaffold. Cell, 2001. 105(1): p. 127–136.

56. Lescar, J., et al., The Fusion Glycoprotein Shell of Semliki Forest Virus: An Icosahedral Assembly Primed for Fusogenic Activation at Endosomal pH. Cell, 2001. 105(1): p. 137–148.

57. Shi, D., A. Sheng, and L. Chi, Glycosaminoglycan-protein interactions and their roles in human disease. Frontiers in Molecular Biosciences, 2021. 8.

58. Tsetsarkin, K.A., C.E. McGee, and S. Higgs, Chikungunya virus adaptation to Aedes albopictus mosquitoes does not correlate with acquisition of cholesterol dependence or decreased pH threshold for fusion reaction. Virol J, 2011. 8: p. 376.

59. Sánchez San Martín, C., et al., Cross-Inhibition of Chikungunya Virus Fusion and Infection by Alphavirus E1 Domain III Proteins. Vol. 87. 2013.

60. Coffey, L.L. and M. Vignuzzi, Host alternation of chikungunya virus increases fitness while restricting population diversity and adaptability to novel selective pressures. Journal of virology, 2011. 85(2): p. 1025–1035.

61. Rückert, C., et al., Impact of simultaneous exposure to arboviruses on infection and transmission by Aedes aegypti mosquitoes. Nat Commun, 2017. 8: p. 15412.

62. Baker, N.A., et al., Electrostatics of nanosystems: application to microtubules and the ribosome. Proceedings of the National Academy of Sciences, 2001. 98(18): p. 10037–10041.

